# AlphaFold models of host-pathogen interactions elucidate the prevalence and structural modes of molecular mimicry

**DOI:** 10.1101/2025.06.04.657796

**Authors:** Delora Baptista, Lidia Gomez-Lucas, Jürgen Jänes, Nevan J. Krogan, Maria João Amorim, Ylva Ivarsson, Pedro Beltrao

## Abstract

Pathogens exploit host cellular machinery through protein-protein interactions (PPIs), often using molecular mimicry to hijack host cellular processes. While there have been thousands of host-pathogen PPIs determined to date, the lack of structural information for these impedes the study of the prevalence of molecular mimicry and convergent evolution of protein interaction interfaces. To address this, we benchmarked AlphaFold2 and 3 for prediction of structures of host-pathogen interactions observing that accurate models can be retrieved when ranking by modelling confidence, despite an overall low performance. We predicted structures for 6,782 pathogen-human PPIs yielding 803 models of higher confidence. Most pathogen proteins interacting with a common human protein are predicted to do so via the same interface, suggesting a high degree of convergent evolution of protein interaction interfaces. When comparing structural models from host-pathogen and host-host interactions, we observe that a majority of pathogen proteins are predicted to target existing human PPI interfaces. We categorized instances of mimicry into different modes, occurring at different frequencies: 1) via the same domain family (least common); 2) via a similar structural motif; and 3) via a similar linear motif (most common). We selected examples of linear motif interactions for binding assay testing, confirming 8 out of 12 predicted interfaces, including 3 viral linear motif interactions. This validates AlphaFold’s ability to model some host-pathogen interactions and the mechanisms underlying molecular mimicry. This work showcases the value of large-scale structural modelling to study convergent evolution of host-pathogen interactions and how molecular mimicry may contribute to infection or host defense.

## Introduction

Many biological processes within a cell are driven by PPIs. During infection, a pathogen must also interact with its host, both to ensure its survival within the host and to support its proliferation. By interacting with host proteins, pathogens can hijack host cellular processes for their own benefit – to enter and exit host cells (in the case of intracellular pathogens), support replication, transport pathogen proteins, evade the host’s immune response, establishing latency or promoting uncontrolled cellular division in oncogenic viruses, among other purposes. To facilitate these host-pathogen PPIs, many pathogens can resort to mimicry of host proteins, either at the sequence or structural level (Elde & Malik, 2009).

Studying molecular mimicry is critical to shed light into the important evolutionary processes that shape host-pathogen interactions. For example, it is largely unknown if pathogens typically evolve to hijack the host by interacting with host proteins through interfaces already used in host PPIs. A structural understanding of molecular mimicry also would be critical for improving drug development pipelines. Molecular mimicry occurs along a spectrum, ranging from local similarity in the form of short linear motifs or the existence of similar structural motifs in small regions of a protein, to more global forms of similarity where entire domains or even the whole protein structure is nearly identical. Global forms of mimicry are more likely to arise through horizontal gene transfer events (Elde & Malik, 2009) which tends to be more common among dsDNA viruses, such as those belonging to Poxviridae and Herpesviridae (Irwin et al., 2021). Known examples of global mimicry of human proteins include viral interleukin-10 homologs in human herpesviruses such as BCRF1 from Epstein-Barr virus (EBV) (Moore et al., 1990), and A38L from vaccinia virus, which is a homolog of human CD47 (Parkinson et al., 1995). More local forms of similarity tend to emerge in pathogen proteins through convergent evolution (Davey et al., 2011; Via et al., 2015). Mimicry of short linear motifs (SLiMs) present in host proteins is extensively used by pathogens (Davey et al., 2011; Sámano-Sánchez & Gibson, 2020; Via et al., 2015), especially viruses (Davey et al., 2011; Mihalič et al., 2023). SLiMs are small, degenerate sequences of 3 to 15 residues, and tend to occur more often in intrinsically disordered regions (IDRs) (M. Kumar et al., 2024). Due to these characteristics, mimicry by SLiMs can be difficult to detect using computational approaches.

Pathogen mimicry can be predicted by sequence similarity searches, but these can be limited in accuracy due to the difficulty in predicting the similarity of interactions and interfaces beyond the most obvious cases of very close sequence similarity of large domains. Mimicry can be uncovered more accurately by comparing the structures of host-pathogen and host-host PPIs. However, the Protein Data Bank (PDB) (Berman, 2000) contains structures for only a very small fraction of all experimentally determined host-pathogen and host-host protein interactions. In addition, these experimental structures do not always cover the full length of the proteins and less structured regions such as IDRs may be more difficult to resolve depending on the experimental method used to obtain the structure. To study the phenomenon of structural mimicry at scale, it may be possible instead to use computational protein structure predictions methods.

AlphaFold2 (AF2) (Jumper et al., 2021), and its multi-chain version, AlphaFold-multimer (AF-multimer) (Evans et al., 2021), have revolutionized protein structure prediction. AF2 and AF-multimer use deep learning to predict protein structure based on the amino acid sequence, relying heavily on coevolutionary information in the form of multiple sequence alignments (MSAs) to infer residue-residue contacts and 3D structure (Jumper et al., 2021; Evans et al., 2021). Host-pathogen PPIs pose additional challenges for alignment-based methods like AF-multimer. There is a lack of coevolutionary information for proteins from different species, and pathogen proteins tend to evolve rapidly, leading to shallower MSAs. Residues at the host-pathogen interface in particular are expected to be under higher selective pressure (Sironi et al., 2015), making it difficult to accurately predict these interfaces. There are other artificial intelligence-based methods for protein structure prediction, such as protein language models that can predict structures based on sequence alone and do not require MSAs (Lin et al., 2023; R. Wu et al., 2022), but these methods tend to perform worse (Elofsson, 2023). AF-multimer has already been used to predict structures for host-pathogen PPIs in several studies (Homma et al., 2023; Bogdanow et al., 2023; Dugied et al., 2025; Kotova et al., 2025), with mixed success. Until recently most of these studies were small and specific, focusing on a single pathogen or even a single protein complex. Finally, AlphaFold3 (AF3) has shown small improvements in the prediction of protein complex structures, which may also translate into higher performance for predictions involving host-pathogen interactions (Abramson et al., 2024).

In this study, we benchmarked AF-multimer and AF3 for the prediction of structures for host-pathogen interactions and attempted to predict the structures of 6,782 interactions, resulting in 803 structures of higher predicted confidence. We compared host-pathogen and human-human interfaces, observing that in the majority of cases the pathogen interacts with existing putative interfaces used by host proteins. The structural models allowed us to classify the host-pathogen interactions into 3 classes of molecular mimicry: domain level, local structured regions, and linear motifs. We predict that the most common form of mimicry is the use of similar SLiMs (from about 4 to 21 amino acids long) located in IDRs, although we also identified several examples of apparent convergent evolution of interfaces at a structural level. Finally, we used peptide binding assays to validate example cases of predicted linear motif interfaces.

Our results show the limitations and the potential for AlphaFold-based predictions of structures of host-pathogen interactions. The predicted structures shed light into the prevalence and structural mechanisms of molecular mimicry driving host-pathogen interactions.

## Results

### Benchmarking AlphaFold models of host-pathogen interactions

To test AF-multimer (Evans et al., 2021) and AF3 (Abramson et al., 2024) performance for the prediction of host-pathogen interactions, we devised a benchmark dataset of 456 experimentally resolved structures of protein pairs (230 viral-mammalian protein pairs and 226 bacterial-mammalian dimers) obtained from the PDB (Berman, 2000) (see Methods and **Table S1**). A total of 452 and 449 protein pairs from the benchmark datasets were successfully modeled with AF-multimer (**Table S2)** and AF3 (**Table S3**), respectively. We evaluated the quality of the predictions by comparing them with the corresponding experimental structures, using MultiMer-align (MM-align) (Mukherjee & Zhang, 2009). Most of the predicted structures have folds that are highly similar to the ones adopted by their experimental counterparts, with most of the models having a template-modeling score (TM-score) (Zhang & Skolnick, 2004) greater than 0.5 (**Fig. 1a**). On the other hand, many of the models have low interface modelling score (DockQ scores, **Fig. 1b**), which suggests that many predicted interfaces for these models have errors. Nevertheless, according to the quality criteria used in the Critical Assessment of PRediction of Interactions (CAPRI) (Lensink et al., 2017), approximately 44.47% (for AF-multimer) and 53.9% (for AF3) of the models would be considered of “Acceptable” quality or higher, indicating that at least the relative positioning of the proteins at the interface is correctly predicted (**Fig. 1c**). We did not observe a significant difference in performance metrics when comparing viral and bacterial protein pairs (**Fig. S1a**). As expected, the performance metrics dropped significantly when considering only protein pairs that were released after the AlphaFold2/3 training set cutoff date (2021-09-30), with 30.11% (for AF-multimer) and 35.16% (for AF3) deemed to be correct (DockQ ≥ 0.23) (**Fig. 1c**). We also confirmed that AF-multimer usually performs worse on pairs with antibodies (**Fig. S1b**), as other authors have noted (Evans et al., 2021; Yin et al., 2022; Yin & Pierce, 2024). These results were expected, considering the presence of antigen-binding hypervariable loops within these proteins.

**Figure 1.**
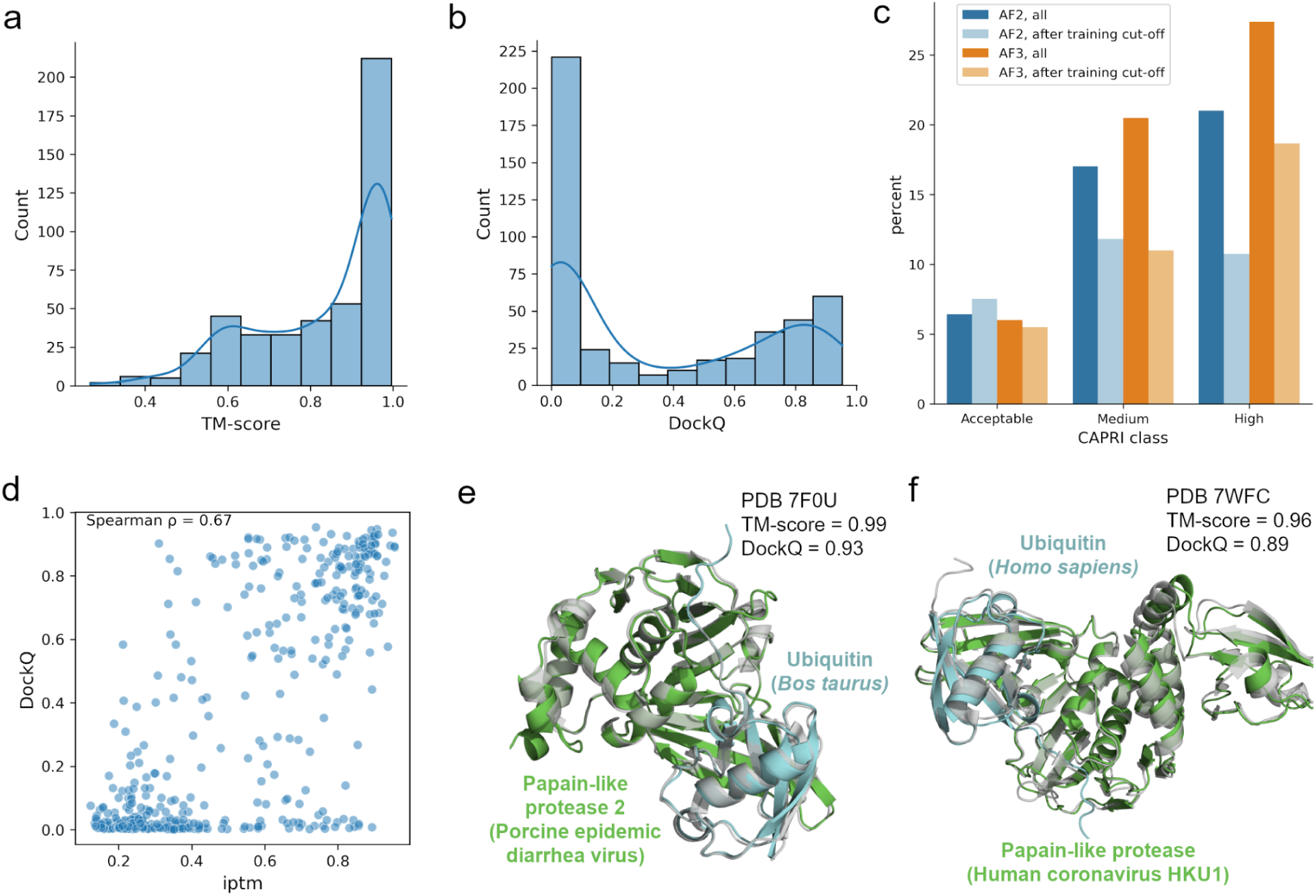
Results for the benchmark datasets. a) Distribution of TM-scores for AF-multimer predictions. b) Distribution of DockQ scores for AF-multimer predictions. c) Number of predicted structures per CAPRI model quality category. Results are shown for structures predicted using either AF-multimer or AF3, and considering all protein pairs in the benchmark dataset or only those with experimental structures released after the AF-multimer v. 2.3.0 and AF3 training data cutoff date (2021-09-30). Models were classified based on their DockQ scores (Acceptable: 0.23 <= DockQ < 0.49; Medium: 0.49 <= DockQ < 0.8; High: DockQ > 0.8). d) Relationship between the ipTM and DockQ scores (Spearman’s ρ = 0.67), for AF-multimer predictions. e, f) Examples of well-modeled structures of viral-human PPIs (colored by chain), superposed onto the corresponding experimental structures from the PDB (in gray). E) Predicted and experimental structures for PDB ID 7F0U, an interaction between papain-like protease 2 (Porcine epidemic diarrhea virus) and ubiquitin (*Bos taurus*). F) Predicted and experimental structures for PDB ID 7WFC, an interaction involving papain-like protease (Human coronavirus HKU1) and ubiquitin (*Homo sapiens*).

We found a positive correlation between the DockQ scores, that measure the overlap between predicted and experimental interfaces, and the interface predicted TM-score (ipTM) interaction modelling confidence estimate scores output by AF-multimer (**Fig. 1d**, Spearman’s ρ=0.67) and AF3 (**Fig. S2b**). This indicates that the ipTM confidence score can be used to select more accurate models. Using an ipTM score of 0.5 as a cutoff to select structures that are more likely to be well-modeled, over 80% of the selected models are of “Acceptable” quality (DockQ ≥ 0.23) or better when using AF-multimer, and over 90% when using AF3. The ipTM scores were more correlated with DockQ scores than pDockQ scores, an alternative interaction model confidence metric (Bryant et al., 2022a; Burke et al., 2023) (**Fig. S1c** and **Fig. S2c**). Finally, the predicted TM-scores (pTM) do not correlate well with the actual TM-score values (**Fig. S1d** and **Fig. S2d**). In light of these results, we decided to use ipTM as the main model confidence metric. These results suggest that, despite the overall relatively poor performance of AlphaFold in modelling host-pathogen interactions, the ipTM score can be used to filter for likely correct models.

We showcase two examples (**Fig. 1e** and **Fig. 1f**) of correctly predicted viral-human PPI structures from our benchmark dataset. In both cases the TM-scores are above 0.9 and the DockQ scores are above 0.8 (models of “High” quality, according to CAPRI criteria), and the figures confirm that the predicted structures (colored by chain) are well aligned with the corresponding experimental structures (in gray).

### Large-scale structural modelling of known host-pathogen interactions

After benchmarking AF-multimer and AF3, we used these to predict the structures of 6,782 high-confidence experimental pathogen-human protein pairs collected from several affinity purification-mass spectrometry (AP-MS) studies and from the Host Pathogen Interactions database (HPIDB) (see Methods and **Table S4**). 6,542 and 6,753 of these pairs were successfully modeled using AF-multimer (**Table S5**) (or an adaptation of AF2, ColabFold (Mirdita et al., 2022), for HPIDB pairs) and AF3 (**Table S6**), respectively, and overall, we observed a general trend for lower ipTM values when using AF3 (**Fig 2a**). Given the modest improvement in the benchmark metrics, the generally lower ipTM values and the restrictive licenses of AF3, we decided to base the rest of our analysis on the results from AF-multimer. Across all datasets, we obtained 803 models with an ipTM score of 0.5 or higher.

**Figure 2.**
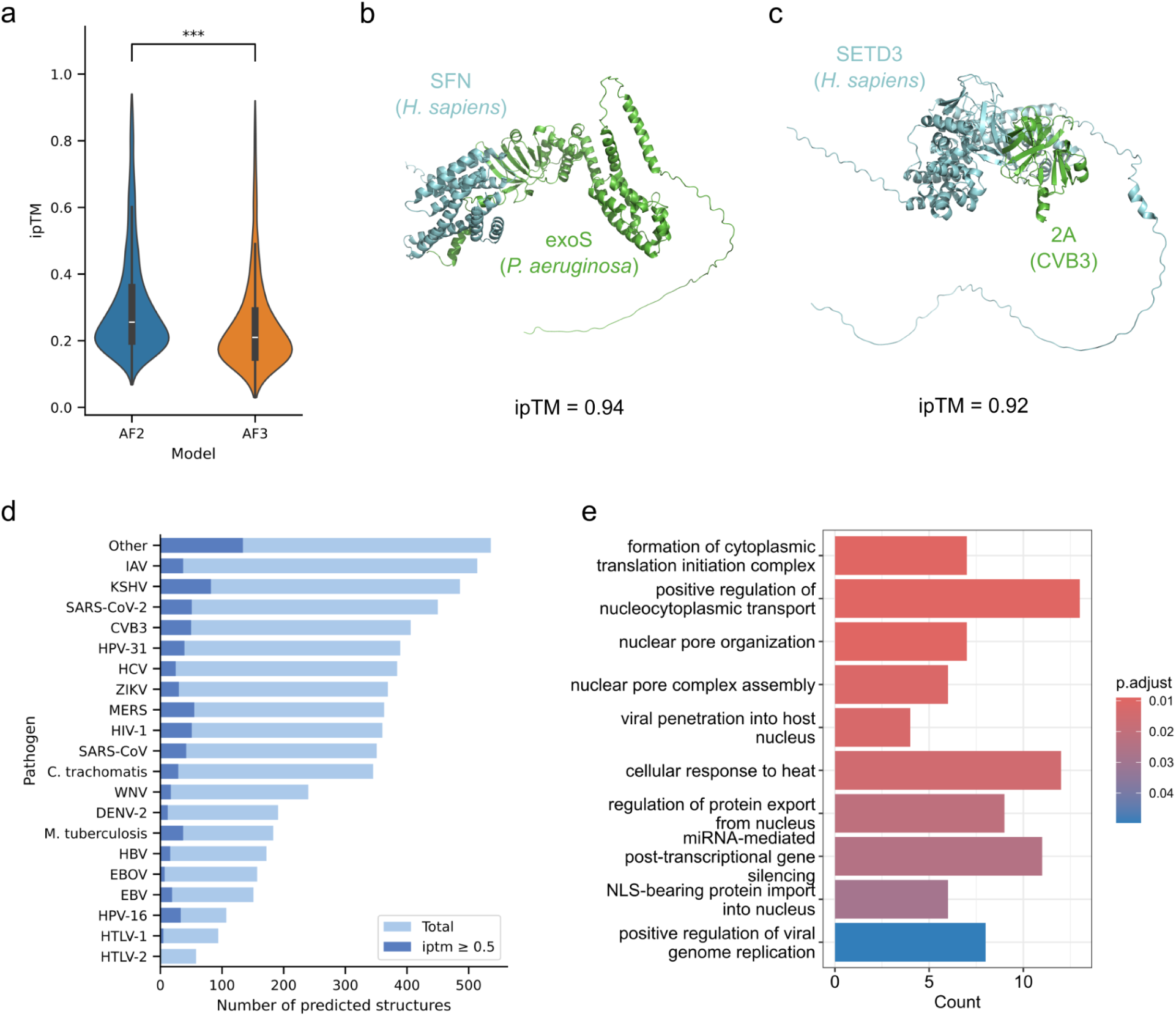
Results for human-pathogen protein pairs from affinity purification-mass spectrometry studies and HPIDB. a) Comparison between AF-multimer and AF3, in terms of ipTM scores. b) AF-multimer (ColabFold) prediction for the interaction between human 14-3-3 protein sigma (SFN), in blue, and Exoenzyme S (exoS) from *Pseudomonas aeruginosa*, in green. c) AF-multimer prediction for the interaction between human protein actin-histidine N-methyltransferase (SETD3), in blue, and protein 2A from Coxsackievirus B3 (CVB3), in green. d) Number of predicted structures (light blue) and structures with an ipTM score >= 0.5 (dark blue), per pathogen. e) GO enrichment analysis results for the human proteins from structures with ipTM >= 0.5.

Two examples of predicted structures with high ipTM scores (> 0.9) are displayed in **Fig. 2b** and **Fig. 2c**. The first is the structure of secreted exoenzyme S (exoS) from *Pseudomonas aeruginosa* interacting with 14-3-3 protein sigma (SFN). The interaction between exoS and a eukaryotic 14-3-3 protein is necessary for the activation of the exoenzyme (Fu et al., 1993), as the 14-3-3 protein acts as a chaperone for exoS (Karlberg et al., 2018). The structure of the homologous interaction between exoS and another 14-3-3 protein is in good agreement with the predicted structure (Karlberg et al., 2018), although it should be noted that this homologous structure was part of the AlphaFold training and template set. Figure **2c** shows the predicted structure of the interaction between protein 2A from Coxsackievirus B3 (CVB3) and human actin-histidine N-methyltransferase (SETD3). SETD3 is a host factor that is essential for enteroviral infection and has been shown to interact with protein 2A (Diep et al., 2019). The structure of 2A in complex with SET3D was recently solved (Peters et al., 2022) and was not used for AlphaFold2/3 training. To exclude the possibility that this experimental structure was used as a template, we repeated the AF-multimer prediction for this interaction excluding template searches. A comparison between the prediction and experimental model showed very good overlap (RMSD=0.97Å), providing further support for the enrichment of correct models at higher confidence estimates.

The number of predicted protein pair structures per pathogen is shown in **Fig. 2d**, illustrating that overall, we did not observe striking differences in the proportion of high confidence models per pathogen. When performing a gene ontology (GO) term enrichment analysis of the identified human partners in these models, we find expected enrichment in processes related with transport into the nucleus and the regulation of transcription and translation (**Fig. 2e**).

Such high confidence models can be used to generate mechanistic and structural hypotheses for future experimental studies. Moreover, the predicted structural models for known host-pathogen interactions can be used to study structural properties related to such interfaces. In the next sections, we focus on studying the targeting of the same human interface by different pathogens and the extent of mimicry when compared to human-human interfaces.

### Pathogen proteins targeting the same interfaces

Some human proteins are known to interact with multiple pathogen proteins. The structural models allow us to predict if the pathogen proteins interacting with the same protein tend to converge on targeting the same interface regions. In our dataset, we identified 90 human proteins (see **Table S9**) having high confidence models (ipTM≥0.5) for 2 or more interacting pathogen proteins, with the top 9 shown in **Fig 3a**. We used the Jaccard index to evaluate the similarity of the interfaces targeted by the same pathogen interactors (scores in **Table S10**), observing that 110 out of 138 pathogen proteins have a Jaccard index ≥ 0.3 (**Fig. 3b**) with at least one other pathogen protein. This result suggests that pathogen proteins most often are predicted to target similar regions in the human proteins they interact with. We illustrate these results with examples that also cover different aspects of pathogens recurrently hijacking the host proteins for different needs, including regulation of trafficking (RAB7A, **Fig. 3c**), nuclear import (importin-α, **Fig. 3d**) and the cell cycle (Protein phosphatase 2A (PP2A), **Fig. 3e**).

**Figure 3.**
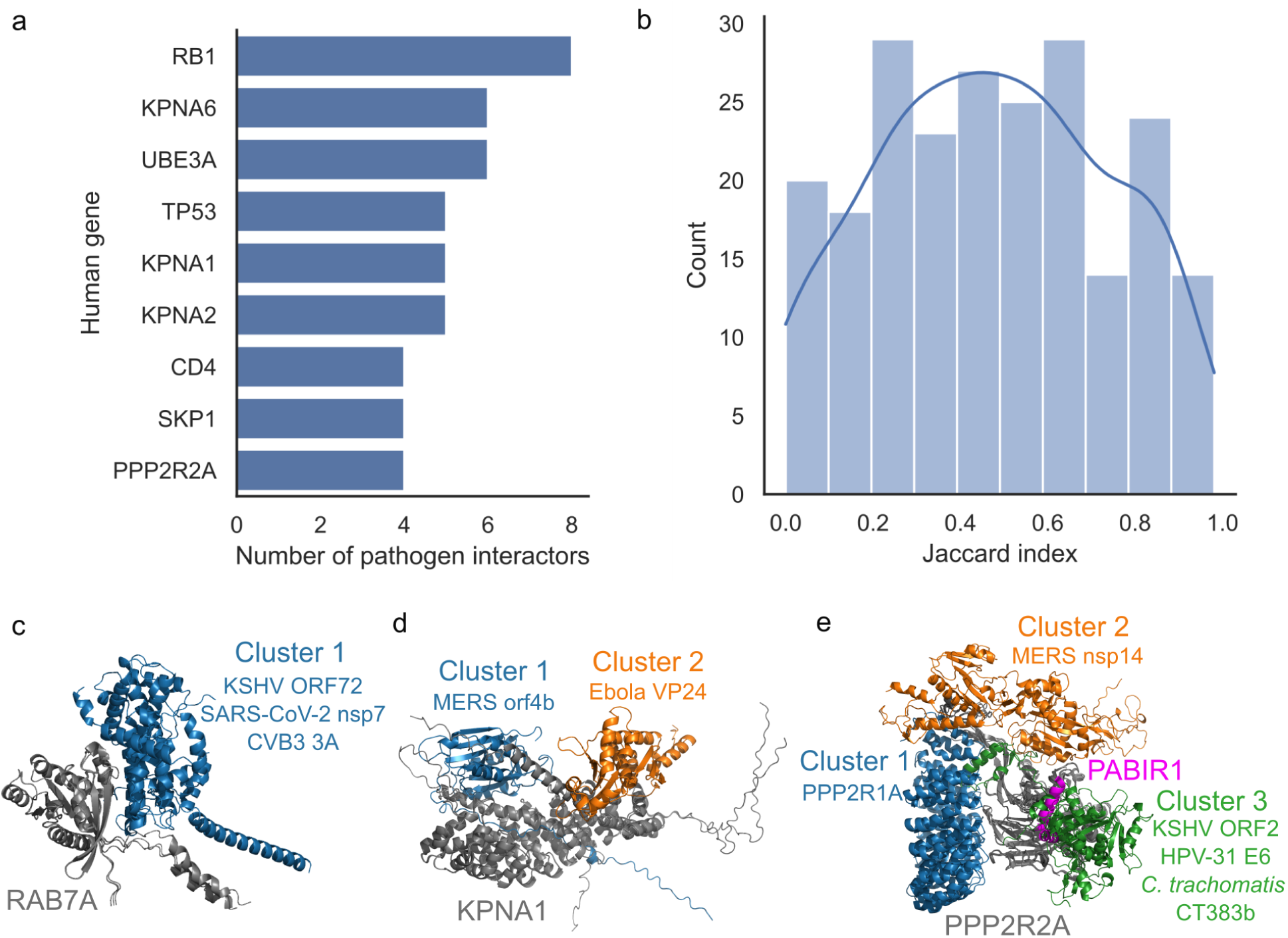
Pathogen interactors targeting the same interface. a) Number of pathogen interactors per human protein (identified by gene symbols). Only human proteins targeted by 4 or more pathogen proteins in our datasets are shown. b) The distribution of Jaccard index values calculated between all pairs of human-pathogen PPIs sharing the same human protein. c), d) and e) Examples of pathogen proteins that target the same interface in the same human protein. c) AF-multimer predicted structures showing viral proteins Kaposi’s sarcoma-associated herpesvirus (KSHV) ORF72, Severe acute respiratory syndrome coronavirus 2 (SARS-CoV-2) nsp7 and CVB3 3A targeting a similar interface in human Ras-related protein Rab-7a (RAB7A). d) AF-multimer structures of Middle east respiratory syndrome coronavirus (MERS) orf4b and Ebola virus VP24 which target different interfaces in importin subunit alpha-5 (KPNA1). For ease of visualization, MERS orf4b was used as the sole representative of cluster 1, which also includes influenza A virus (IAV) nucleoprotein, EBV EBNA1, and KSHV ORF17. e) AF-multimer structures of KSHV ORF2, MERS nsp14, Human papillomavirus 31 (HPV-31) E6, *Chlamydia trachomatis* CT383b (residues 156-243) and human protein PPP2R1A interacting with PPP2R2A. The structures were additionally superposed onto an experimental structure (PDB 8SO0 (Padi et al., 2024)) of PPP2R2A interacting with PABIR1.

Ras-related protein Rab-7a (RAB7A) plays an important role in endolysosomal trafficking (Feng et al., 1995; Bucci et al., 2000) and has been shown to be involved in viral and bacterial infections across many pathogens (Caillet et al., 2011; Daniloski et al., 2021; D’Costa et al., 2015; Macovei et al., 2013). RAB7A is a GTPase, active when in its guanosine triphosphate (GTP)-bound form. Among our predicted structures (ipTM >= 0.5), RAB7A is found in models with 3 different pathogen proteins – CVB3 3A, Severe acute respiratory syndrome coronavirus 2 (SARS-CoV-2) nsp7, and Kaposi’s sarcoma-associated herpesvirus (KSHV) ORF72. All 3 proteins bind to a highly similar interface in RAB7A (**Fig. 3c**, high Jaccard Index values), despite not being structurally similar. In all 3 cases, the viral proteins do not seem to interfere with the GTP-binding site in RAB7A, since only one of the GTP-binding site residues (residue 40) is in contact with the viral proteins. When comparing the predicted structures with an experimentally determined structure of RAB7A bound to GTP (PDB 1T91 (M. Wu et al., 2005)), we find that the predicted RAB7A structure adopts a conformation that is similar to its GTP-bound form when interacting with 3A or ORF72, but it assumes a slightly different conformation when interacting with nsp7, possibly suggesting that different viral proteins might bind to different (GTP- or guanosine diphosphate (GDP)-bound) forms of RAB7A (**Fig. S4**). The RAB7A interface targeted by the viral proteins is also similar to the interface used by human proteins that interact with RAB7A (**Fig. S5**, same cluster). These pathogen proteins are from unrelated viral families belonging to Picornaviridae, Coronaviridae and Herpesviridae, with positive and negative sense RNA, and DNA genomes, respectively. The data reflects how important RAB7A is for infection of phylogenetically different viruses and the degree of likely convergent evolution that can occur in targeting the same host interface region.

Many viral and host proteins need to access the nucleus for specific functions. Importin-α proteins are involved in the transport of proteins into the nucleus, recognizing nuclear localization signals (NLS) in the proteins to be translocated (Moroianu et al., 1995). During infection, viral proteins interact with importin-α in order to be transported into the nucleus or to prevent the nuclear import of host proteins involved in host cell responses including innate immunity activation (Vogel et al., 2023). Among our high confidence models, we have predicted structures for interactions between importin subunit alpha-5 (KPNA1) and 5 viral proteins of unrelated viral families - Influenza A virus (IAV) H1N1 nucleoprotein (NP, P03466), Epstein-Barr nuclear antigen 1 (EBNA1, P03211), Ebola virus VP24, KSHV ORF17 and Middle East respiratory syndrome coronavirus (MERS) orf4b. All of the viral proteins that interact with KPNA1 use a similar interface, except VP24 (**Fig. 3d**). When clustering the interactions based on the interface residues in KPNA1 that are targeted by each interactor (**Fig. S6**), we find that the interface used by VP24 is more similar to the interface used by human proteins importin subunit beta-1 (Q14974) and importin-5 (O00410). The main difference between the interface used by VP24 and the interface targeted by other viral proteins is the lack of interaction with residues within the major NLS binding site in KPNA1 (residues 149-241), although it does interact with a few residues within the minor NLS binding site (318-406), unlike the other interactors within the same cluster. Consistent with these predictions, all of these viral protein interactors can translocate to the nucleus during infection (Digard et al., 1999; Matthews et al., 2014; Petti et al., 1990; Tsurumi et al., 2021; Vogel et al., 2024), although VP24 is preferentially located in the cytoplasm (Han et al., 2003; Nanbo et al., 2013; Vogel et al., 2024). VP24 is also known to inhibit the interferon-induced nuclear import of tyrosine-phosphorylated STAT1 (PY-STAT1) by interacting with the PY-STAT1 binding region in KPNA1, located near its C-terminal (Reid et al., 2007).

Intracellular pathogens frequently manipulate the host’s cell cycle for their own benefit (Fan et al., 2018). PP2A is a multi-subunit protein that plays an important role in regulating the cell cycle (Wlodarchak & Xing, 2016), making it an attractive target for pathogens. In our models (ipTM >= 0.5), the serine/threonine-protein phosphatase 2A 55 kDa regulatory subunit B alpha isoform (PPP2R2A) has 3 different viral protein interactors (KSHV ORF2, MERS nsp14, and HPV-31 E6) and a bacterial interactor (*Chlamydia trachomatis* CT383b-156-243). Interestingly, the interface in PPP2R2A targeted by CT383b-156-243 is similar to the one used by ORF2 and E6, while MERS nsp14 is predicted to bind to a different interface. Fig. 3E shows the structures of the host-pathogen pairs superposed onto an experimental structure (PDB 8SO0 (Padi et al., 2024)) of PPP2R2A interacting with PPP2R1A-PPP2R2A-interacting phosphatase regulator 1 (PABIR1), an inhibitor of PP2A. CT383b-156-243, ORF2 and E6 interact with PPP2R2A and are predicted to bind the same region as PABIR1. A different PP2A inhibitor, alpha-endosulfine (Uniprot ID O43768) also binds to PPP2R2A using a similar interface (**Fig. S7**). These similarities could help explain how these pathogen proteins might use this interaction to induce cell cycle arrest. MERS nsp14 neither uses the PABIR1 binding site, nor the interfaces where the other PP2A subunits interact with PPP2R2A. Instead, it targets an interface in PPP2R2A that is similar to the one used by the human large ribosomal subunit protein uL11 (RPL12, Uniprot ID P30050) (**Fig. S7**).

Our results indicate that, when different pathogen proteins interact with the same host protein, they will most often be predicted to target the same interface. Given that the interactions are derived from widely diverse pathogens, this shows that convergent evolution at the level of PPIs is likely to be extremely prevalent.

### Pathogen mimicry of human-human interfaces

In several of the examples above, we noted that the pathogen proteins used similar interfaces as those used by existing human PPIs. These are potential examples of mimicry where proteins from pathogens copy the structural properties of human interactions. We compared the host-pathogen interaction models with a previously generated dataset of predicted structures of human interactions, covering over 14,549 human proteins (Burke et al., 2023; Jänes et al., 2024). We first determined if the pathogen proteins occupy the same interface used in a human protein interaction (scores in **Table S11**). In 407 out of 803 human-pathogen PPIs, the pathogen protein targets an interface that is at least partially similar to a predicted interface targeted by human proteins (Jaccard index of the targeted interfaces ≥ 0.3. In addition, we used iAlign to see if the interfaces of pathogen-human PPIs and human-human PPIs involving the same shared protein are structurally similar. 178 human-pathogen PPIs have an iAlign score of 0.5 or higher when compared with human-human PPIs, suggesting a potential convergence to a similar structure at the interface. We manually inspected these cases and identified 94 of these host-pathogen interfaces having clearer structural similarity to human-human interfaces.

We then attempted to classify interface similarity into three main categories: 1) same domain family; 2) similar local structured motif; 3) similar linear peptide motif (**Fig. 4**, **Table S12**). In the first case, the pathogen-human and human-human interfaces are similar because the pathogen protein interacts via a domain that belongs to the same or a highly similar domain family (high iAlign iTM-score values). In the example provided in **Fig. 4b**, both the pathogen protein (ORF72, a viral cyclin homolog from KSHV and the human protein (G1/S-specific cyclin-D2) that interact with the shared protein, human cyclin dependent kinase 2 (CDK2), belong to the same family. When superposed, the structures are well aligned, showing that the similarity extends beyond the interface region. Such cases of interaction mimicry are unlikely to occur by convergent evolution and are more plausibly explained by horizontal gene transfer.

**Figure 4.**
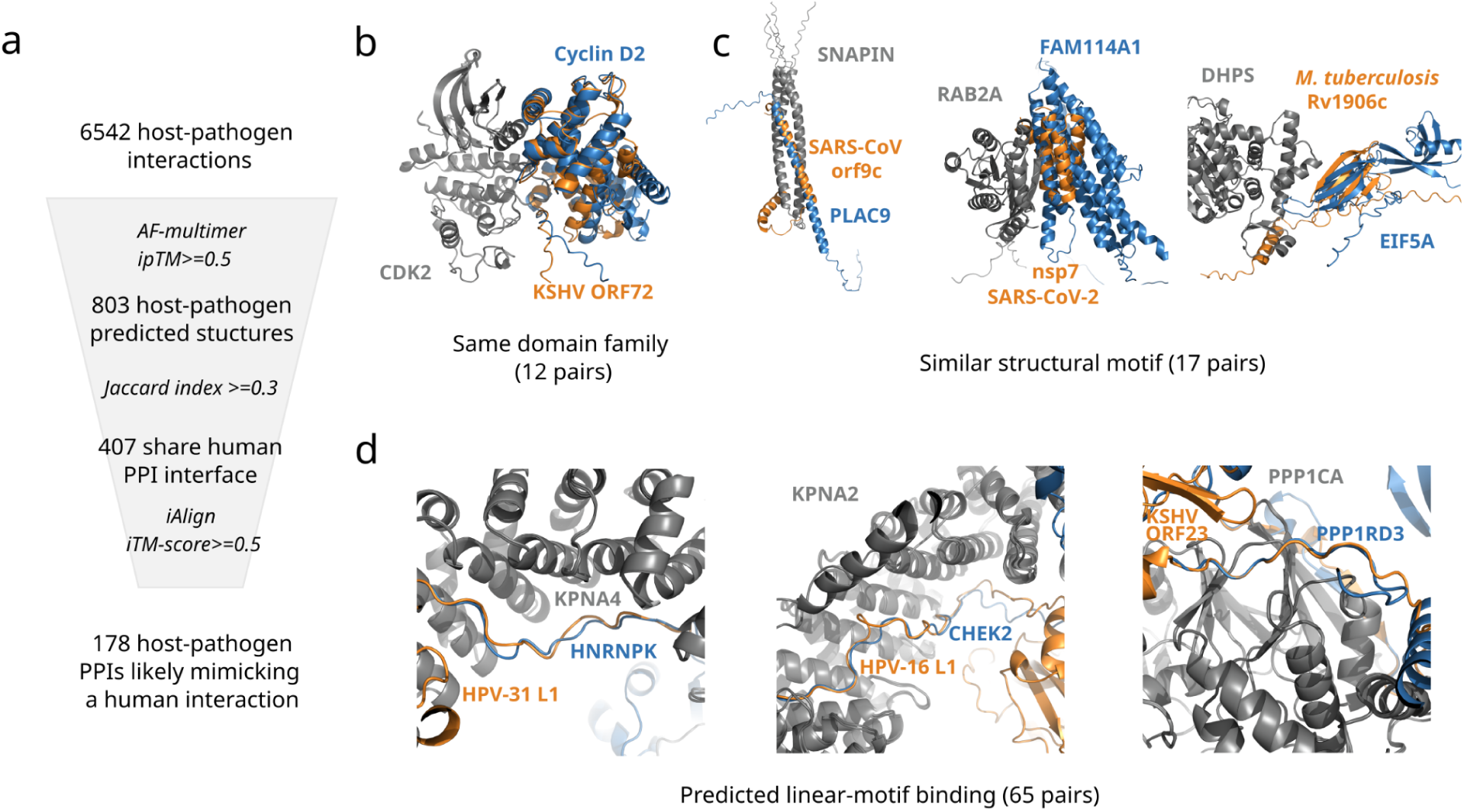
Structural modes of host-pathogen mimicry from predicted structural models. a) The host-pathogen models of higher confidence were compared with a dataset of 37,344 predicted structures for human protein interactions to identify host-pathogen interfaces similar to human-human interfaces. A set of 178 host-pathogen models were predicted as likely cases of mimicry that were manually inspected to identify those that are similar due to: b) belonging to a similar domain family; c) a similar structural motif without having a similar domain family; d) mostly linear peptide interactions often sharing a few interacting residues - predicted linear-motif interactions.

Only 12 instances of mimicry using a similar domain family were easily identified. For the majority of host-pathogen PPIs, however, when structural similarity exists, it is usually limited to the interface and neighboring residues. In these cases, the pathogen protein interfaces contain structural motifs that bear a resemblance to the structural motifs found in the interfaces of human proteins, but these interfaces do not belong to the same type of domain families. These cases constitute potential examples of convergent evolution of small structured interface regions. Three examples of host-pathogen interactions with this type of interface similarity are shown in **Fig. 4c**.

In the first example we show that the structure of the interface that the Severe acute respiratory syndrome coronavirus (SARS-CoV-1) orf9c (previously called orf14) protein uses to interact with SNARE-associated protein Snapin (SNAPIN) is highly similar (iAlign iTM-score=0.75) to that of human Placenta-specific protein 9 (PLAC9). SNAPIN is a member of the biogenesis of lysosome-related organelles complex 1 (BLOC-1) (Starcevic & Dell’Angelica, 2004), and it is also part of the BLOC-one-related complex (BORC), which regulates the positioning and movement of lysosomes (Pu et al., 2015). Since β-coronaviruses use lysosomes for egress (Ghosh et al., 2020), orf9c might have evolved to mimic the interface structure of human proteins that interact with SNAPIN so that it could hijack it and manipulate lysosomal trafficking.

The second example displays the SARS-CoV-2 nsp7 and human protein NOXP20 (FAM114A1) interacting with Ras-related protein Rab-2A (RAB2A), a human protein that regulates endoplasmic reticulum-Golgi trafficking (Kajiho et al., 2016). FAM114A1 and nsp7 target a similar interface in RAB2A (Jaccard index=0.63). The proteins have a structurally similar interface (iAlign iTM-score = 0.61), both having helices occupying similar positions near the interface. We hypothesize that nsp7 adopts a structure at the interface similar to that of human proteins that interact with RAB2A to direct host membrane trafficking processes towards the trafficking of viral proteins.

In the final example, we show that protein Rv1906c from *Mycobacterium tuberculosis* and isoform 2 of human eukaryotic translation initiation factor 5A-1 (EIF5A) are predicted to use a similar structural motif (beta-strands) to interact with deoxyhypusine synthase (DHPS). The beta-strands occupy similar (but not the same) positions relative to DHPS and the residues they interact with in DHPS are partly the same (Jaccard index = 0.54). DHPS is involved in hypusination, a post-translational modification unique to EIF5A in which a lysine residue is modified to become a hypusine residue (Ganapathi et al., 2019). It has been shown that some bacterial infections can increase hypusination of EIF5A in macrophages, as part of the antimicrobial response of the host (Gobert et al., 2020). Since *Mycobacterium tuberculosis* infects macrophages (Bussi & Gutierrez, 2019), it appears that the structure of Rv1906c may have evolved to mimic the part of the structure of EIF5A that binds to DHPS, potentially blocking the EIF5A-DHPS interaction and inhibiting the host’s immune response.

The most common (65 out 94 manually inspected cases) of predicted mechanism of interaction mimicry was observed for interactions using short linear motifs (**Fig. 4c**). In these cases, at least a part of the pathogen interface is linear/less structured, similar to the interfaces of human proteins that interact with the same target proteins. Among these host-pathogen pairs, we found examples where the interfaces of the pathogen proteins contain short linear motifs that are also present in the interfaces of other binding partners. Six examples are shown in **Fig. 5**, that we describe in more detail below. A more extensive list of examples of pathogen mimicry of human short linear motifs found in our datasets is provided in **Table S13**. Additionally, we note that we also found 29 pairs where the pathogen protein targets an interface not seen in predicted structures of human-human PPIs (Jaccard Index = 0), suggesting that these could potentially contain novel binding interfaces exploited by pathogens (see **Table S14**).

**Figure 5.**
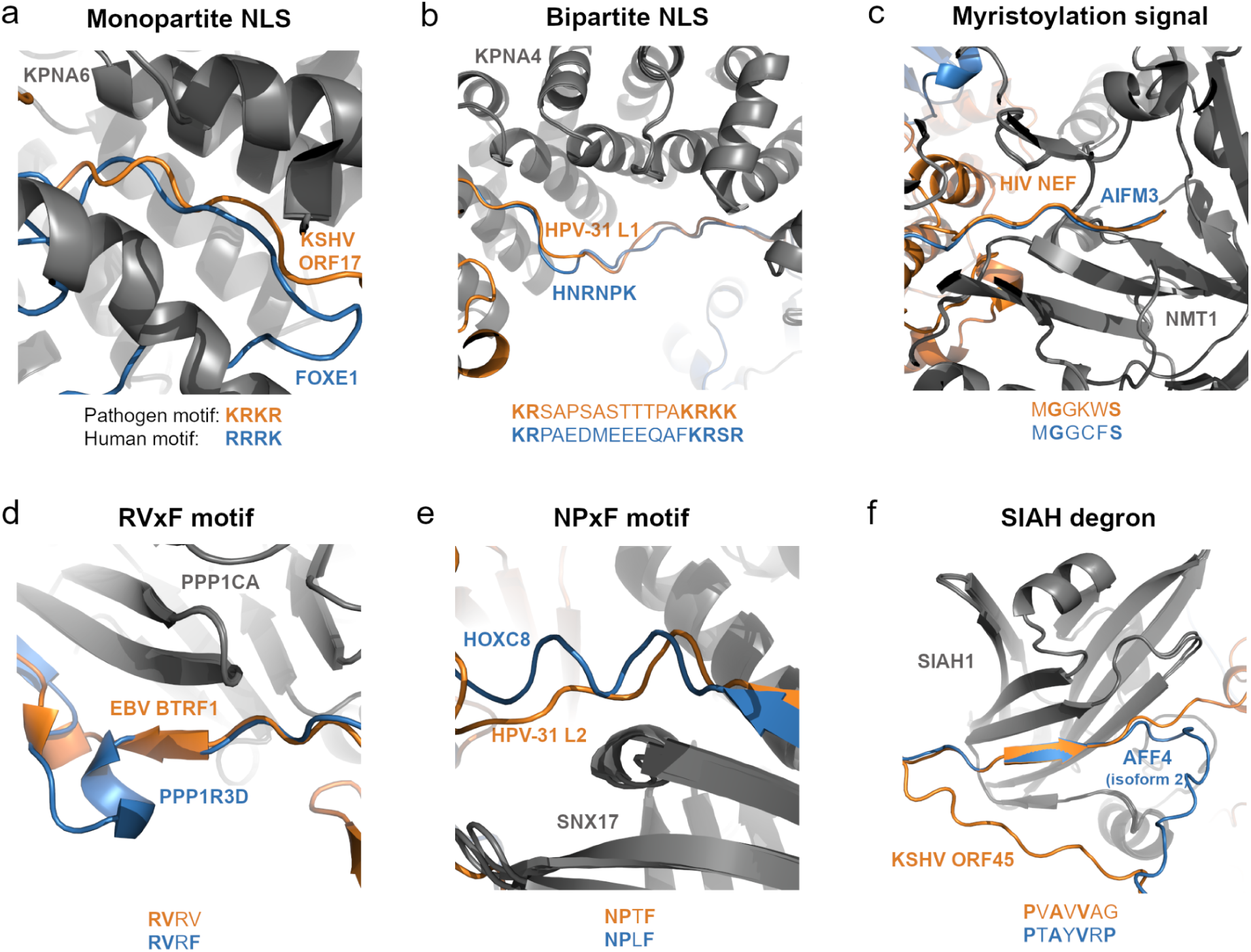
Examples of human-pathogen interfaces that are structurally similar to human-human interfaces and contain similar short linear motifs. Pathogen proteins and linear motifs are shown in orange, human interactors are shown in blue, and the shared proteins are dark gray. a) Monopartite NLSs present in KSHV ORF17 and human protein FOXE1. b) Bipartite NLSs present in HPV-31 L1 and human protein HNRNPK. c) Myristoylation signals in HIV NEF and AIFM3. d) A partial RVxF motif present in KSHV ORF23 and a full RVxF motif found in human protein PPP1R3D. e) NPxF motifs found in HPV-31 L2 and human protein HOXC8. f) A partial SIAH degron (PxAxVxP motif) in KSHV ORF45 and a full SIAH degron present in isoform 2 of human protein AFF4.

### Examples of pathogen mimicry of human interfaces via linear-motif interactions

Many of the examples of mimicry by linear motifs that we predicted are interactions between pathogen proteins that carry out functions in the nucleus and human proteins involved in nuclear transport (importins). As expected, we find that these pathogen proteins have linear interfaces that contain either monopartite or bipartite NLSs (Lu et al., 2021) recognized by importin-α proteins. **Figure 5a** is an example of a monopartite NLS found in a disordered region of the KSHV capsid scaffolding protein (ORF17). This NLS is recognized by the same importin subunit alpha-7 (KPNA6) that recognizes a similar motif present in human forkhead box protein E1 (FOXE1). **Figure 5b** shows the interaction between HPV-31 L1 and importin subunit alpha-3 (KPNA4). L1 has a bipartite NLS motif that occupies the same position near the interface as a NLS present in human heterogeneous nuclear ribonucleoprotein K (HNRNPK), which also interacts with KPNA4.

In addition to NLS motifs, we also identified other motifs in pathogen proteins with linear interfaces. A small disordered region of the human immunodeficiency virus 1 (HIV) Nef protein has an N-terminal myristoylation signal (GxxxS motif), similar to the interface of Apoptosis-inducing factor 3 (AIFM3) (**Fig. 5c**). The GxxxS motif is recognized by glycylpeptide N-tetradecanoyltransferase 1 (NMT1), which adds a myristoyl group to the N-terminal glycine residue of this motif (Resh, 1999). N-myristoylation of Nef localizes it to cellular membranes (Yu & Felsted, 1992).

The BTRF1 protein from EBV has a disordered interface region bearing a partial RVxF motif which is recognized by several members of the protein phosphatase 1 (PP1) catalytic subunit family of proteins. **Fig. 5d** shows the interface region of the interaction between BTRF1 and serine/threonine-protein phosphatase PP1-alpha catalytic subunit (PPP1CA), as well as the interface of the interaction between PPP1CA and the human protein phosphatase 1 regulatory subunit 3D (PPP1R3D). The interface of PPP1R3D has an RVxF motif in the same position as the partial RVxF motif (“RVR”) of BTRF1. The RVxF binding site of PPP1CA is a non-catalytic binding site. We hypothesize that BTRF1 mimics the RVxF containing interfaces of PP1 regulatory subunits to manipulate PP1 activity for the virus’ own benefit. An analogous protein in KSHV, ORF23, also interacts with PPP1CA (Davis et al., 2015), and it has been proposed that the interaction between ORF23 and PPP1CA might promote the dephosphorylation of RNA polymerase II, leading to its recruitment to a virus-specific transcription preinitiation complex (Nishimura et al., 2017). The interaction between BTRF1 and PPP1CA might serve a similar purpose.

Viruses have evolved many sophisticated mechanisms to enter cells. Many use endocytosis to access the cellular environment, from which they need to escape. The HPV L2 protein assists in the escape of viral DNA from endosomal compartments through interaction with sorting nexin-17 (SNX17) (Bergant Marušič et al., 2012), which regulates endosomal recycling of cell surface proteins (Knauth et al., 2005; Steinberg et al., 2012). SNX17 recognizes its targets based on an NPxY/F motif (Stiegler et al., 2014). In **Fig. 5e**, we show that in our AF-multimer predicted structure SNX17 binds to an NPxF motif (NPTF) present within a disordered region in HPV-31 L2, and that HOXC8, the human SNX17 interactor with the most structurally similar interface, also has an NPxF motif at the interface (NPLF). The structure and sequence of HPV-31 L2 seem to have evolved to mimic the interfaces of human proteins that interact with SNX17, allowing it to compete with these human proteins for SNX17. However, it has been proposed that the NPxF/Y motif bound by SNX1 in HPV-31 L2 is ^249^NPAY (Bergant Marušič et al., 2012) or the extended SNX17 binding motif ^247^YENPAY (F/YxNPxF/Y) (Bergant & Banks, 2013), instead of the ^155^NPTF motif identified in our models. For HPV-16, a different papillomavirus, experimental evidence seems to indicate that the F/YxNPxF/Y motif is essential for the interaction between L2 and SNX17, while the corresponding NPTF motif is not (Bergant & Banks, 2013), although a different study suggests that HPV-16 L2 may have additional SNX17 binding sites (Martín-González et al., 2024). Since all of our AF-multimer and AF3 models predicted that the ^155^NPTF motif was located at the interface, and since the ipTM score of the best AF-multimer structure is 0.73, we decided to test this interaction experimentally via a peptide binding assay (see the subsection below).

KSHV has a lytic and a latent cycle, and KSHV’s ORF45 is a virion-associated tegument protein expressed during lytic infection, being an immediate-early gene, responsible for diverse functions from early to late phases in infection. ORF45 and ALF Transcription Elongation Factor 4 (AFF4) both interact with E3 ubiquitin-protein ligase SIAH1, and are predicted to interact via a similar interface (Jaccard index = 0.64), with both proteins having a less structured region near the interface (**Fig. 5f**). SIAH1 targets proteins for ubiquitination and proteasome-mediated degradation, and its substrates usually contain a SIAH degron, PxAxVxP (House et al., 2003). This motif is present in the predicted interface region of AFF4 (PTAYVRP). The ORF45 interface also has a partial SIAH binding motif (PVAVVAG), located within the region where the pathogen interface mimics the human interactor’s interface. This prediction is in line with the observation that the interaction of SIAH1 with ORF45 and its targeting for degradation depend on residues of ORF45 that are predicted to be at the interface (Abada et al., 2008).

### Peptide binding assays for predicted linear-motif interactions

Our analysis pinpointed specific linear-peptide regions that are predicted to be important for host-pathogen interactions, most of these not being previously known. From the above described examples, we selected 3 human proteins (SIAH, PPP1CA and SNX17) involved in predicted linear motif host-pathogen interactions. Based on the human-human and pathogen-human interaction models, we selected 12 peptides (6 viral and 6 human) that were predicted to be at the interface of one of these proteins, together with 3 putative positive control peptides, and subjected these to peptide binding assays (**Fig. 6**). The positive controls confirmed binding for SIAH1 and SNX17 but not for PPP1CA, although we did observe binding peptides for PPP1CA among our predictions. Out of the 12 predicted interactions we confirmed 8 (66%) but with a higher success rate observed for the human-human interactions (5 out of 6) than for the host-pathogen interactions (3 out of 6). For the viral proteins, we confirmed binding between PPP1CA and peptides in ICP34.5 of HHV-1 (TPARVRFSPHVRVRH) and BTRF1 of HHV-4 (SVKRVRVDEGANTRR) as well as binding between SIAH1 and ORF45 KSHV (TPKPVAVVAGRVRSS). From these 3 validated examples, there is additional experimental evidence supporting the correct identification of the interface region for the interactions between SIAH1 and ORF45 and for PPP1CA and ICP34.5 (Y. Li et al., 2011). For the SNX17 4.1/ezrin/radixin/moesin (FERM)-like domain, which recognizes NPxY/F motifs (Stiegler et al., 2014), we validated a predicted interaction with a NPxF motif of the human homeobox protein Hox-C8 (HOXC8), but failed to validate the predicted NPxF-dependent interaction with the HPV-31 L2 protein, in line with previous literature characterizing the interaction between SNX17 and L2 from a different human papillomavirus, HPV-16 (Bergant & Banks, 2013). The observed interactions were found to be mostly specific, with the major exception being that PPP1CA also bound several of the predicted SIAH1 target peptides.

**Figure 6.**
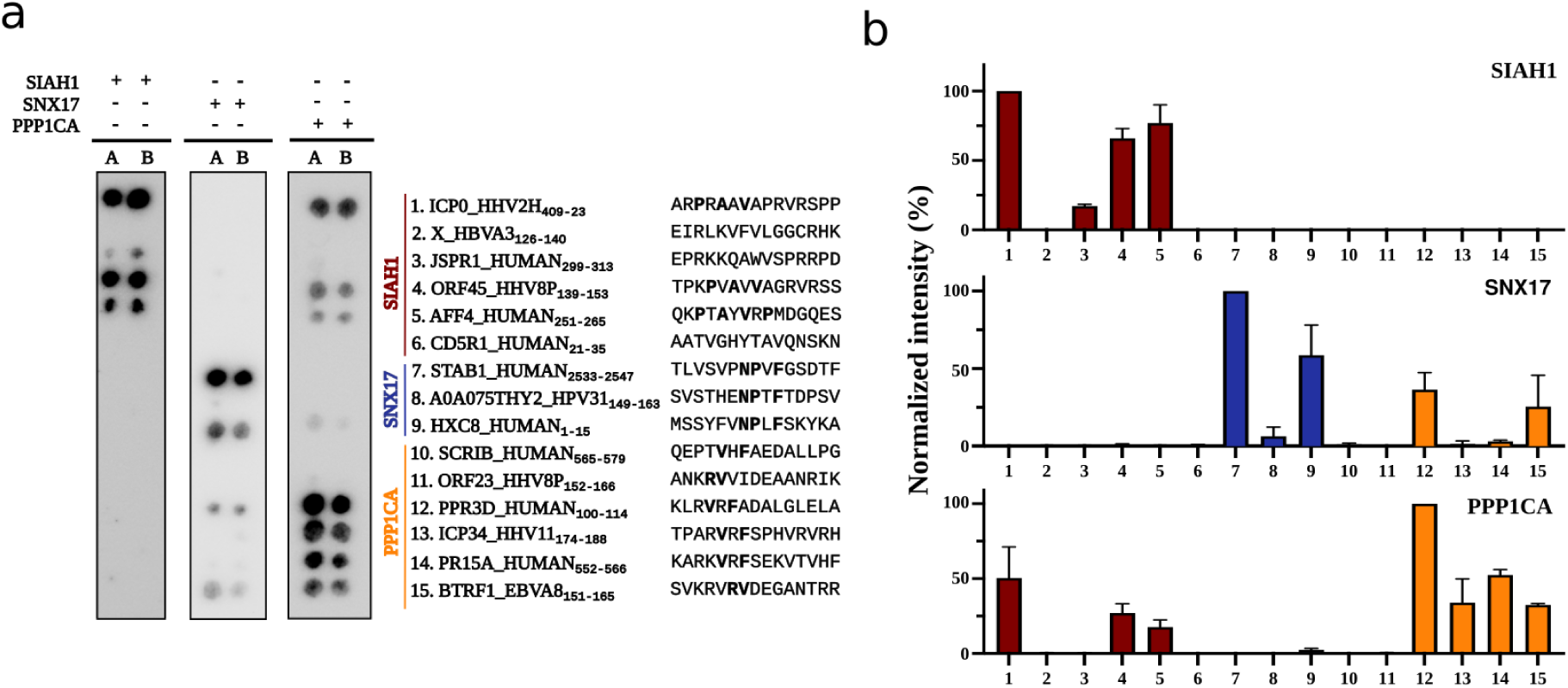
Binding specificity of proteins assessed by SPOT arrays. a) Membranes containing immobilized human and viral peptides predicted to bind to SIAH1, SNX17 or PPP1CA were incubated with the corresponding GST-tagged proteins. The array designs also included previously found interactors to serve as positive controls (peptide 1, 7 and 10 (Adachi & Tsujimoto, 2010; Nagel et al., 2011). The peptide sequences are shown on the right. b) Quantification of normalized signal for each peptide spot. Bars represent mean values ± SD from two replicates (n=2)

For the interaction between SIAH1 and ORF45 KSHV (TPKPVAVVAGRVRSS), we also performed an alanine scanning experiment where each position was mutated to alanine to measure the impact of affecting each position in the peptide binding assay (**Fig. S8**). The strongest observed effect was the mutation of the valine at position 8 which is part of the putative partial match to the SIAH binding motif PxAxVxP. This valine residue has also been previously shown to be important for SIAH binding to ORF45 *in vivo* (Abada et al., 2008) This confirms not just the binding to the peptide, but also the importance of this valine residue in making contacts at the interface as predicted by the structural model.

These results suggest that the models generated by AF-multimer can be used to prioritize likely binding sites, while at the same time indicating that there might be an increased error rate in the validation of interaction regions predicted for host-pathogen interactions.

## Discussion

Molecular mimicry of host proteins is a strategy commonly used by pathogens to hijack host cellular processes. As mimicry of protein interfaces is difficult to predict from sequence alone, the extent this occurs is largely uncharacterised. When detailed, it may provide evidence of pathogenesis mechanisms and expose previously unrecognised regulatory functions of host proteins. In this study, we used AlphaFold to model the structures of thousands of host-pathogen PPIs, obtaining a small but sizable set of higher-quality predictions. We then compared these structures with structural models of human-human PPIs to try to uncover previously uncharacterized examples of molecular mimicry and to better understand this phenomenon at the structural level. We found that pathogen and human proteins targeting the same human protein often bind to similar regions in the shared protein. Pathogens may adopt this strategy to redirect the targeted protein towards processes essential for their own life cycles or as a way to disrupt and inhibit human-human PPIs by blocking access to the shared protein.

We had previously performed large-scale AlphaFold modelling of human PPIs, noting that, for a given protein, most interaction partners are predicted to occupy the same interface (Jänes et al., 2024). The biological interpretation for this would be that many interactions in cells are mutually exclusive. However, a technical limitation may be that AlphaFold may be biased to preferentially build models to a specific interface region due to signals captured in the multiple sequence alignments. In addition, it is possible that AlphaFold may report higher confidence for models that resemble those in the training data, even if the similarity is local. For these reasons, the degree of interaction mimicry predicted here should be considered an upper bound.

Linear motif interactions have often been described as a key structural mechanism for molecular mimicry employed by pathogens (Davey et al., 2011; Via et al., 2015). Our structural analysis allowed us to classify interaction mimicry by 3 structural classes, suggesting that the majority of pathogen proteins that employ molecular mimicry do so by this mechanism. To a lesser extent, we also identified examples of structural mimicry occurring only at the interface within folded regions, possibly acquired by the pathogen protein through convergent evolution. We provide evidence from the existing literature which supports the existence of the interfaces and SLiMs we have identified and their relevance for infection. In addition, we were able to experimentally validate three viral linear motif interfaces, providing further proof that AlphaFold can accurately model at least some host-pathogen interfaces and that a similar method to the one described in this study could be applied to uncover instances of molecular mimicry in other host-pathogen PPI datasets. However, the lower success rate in the experimental validation of the predicted host-pathogen interactions (50%) compared with human-human interactions further highlights the need for experimental validation, in particular in the context of host-pathogen interactions.

We observed cases where pathogens manage to interact with human proteins without resorting to mimicry, such as when a pathogen protein that interacts with a known interface in the host protein lacks structural similarity with other interactors. In addition, we identified several examples where a pathogen protein targets an entirely different interface, not seen among our set of human-human PPIs. Additional experimental evidence would be required to confirm that these novel interfaces in our structural models are correct. Such cases could be particularly relevant for the development of therapies as disrupting such interactions would less likely disrupt the normal function of the human proteins. The identification of a subset of host targets that have multiple pathogen interactors all binding to a similar interface may also facilitate the discovery of pan-pathogen antimicrobials.

In the cell, many pathogen and human proteins exist as part of complexes and not as single proteins. In addition, some of these proteins might also interact with nucleic acids. Here we only modeled host-pathogen protein pairs, since obtaining stoichiometric information for larger protein complexes is not possible. Since the predicted interface residues might not be accessible when the proteins are part of larger complexes, this could have led to the misidentification of certain interfaces. This approach might also fail to identify interfaces that only exist in multimeric forms of a given protein. In the future it would be interesting to extend our analysis to higher-order complexes, to determine how adding more biomolecules would affect interface prediction. Analyzing the conformational changes induced by post-translational modifications and using methods to explore the full conformational space of the structures would also be of interest.

Two recent studies have also focused on the structural prediction of host-pathogen interactions. Saluri et al. predicted the structures of over 8,000 pathogen-human PPI pairs from HPIDB (Saluri et al., 2024) using FoldDock (Bryant et al., 2022b), an adaptation of AF2. In the second study, Li and colleagues benchmarked several protein structure prediction methods and then used AF-multimer 2.2.0 to predict the structures of more than 11,000 viral-human protein pairs (L. Li et al., 2025). Like our study, both showed that it is possible to predict structures of host-pathogen interactions, although Saluri and colleagues also emphasized that host-pathogen interactions tend to be more difficult to model than human-human interactions, a tendency that was also observed in our study. Li and colleagues showed that predicted interface regions have differential evolutionary patterns, in line with their importance for host-pathogen interactions. They also discuss how some host-pathogen interactions overlap with known human interfaces. Our work goes beyond these studies by focusing on the structural mechanisms of mimicry (e.g. full domain, folded regions, linear motifs).

In conclusion, predicted structural models of host-pathogen interactions provide insight into the strategies used by different pathogens to hijack host cellular processes to promote their replication and survival within the host, which may be useful for the development of novel therapies.

## Methods

### Benchmarking datasets

We searched the Research Collaboratory for Structural Bioinformatics (RCSB) PDB (Berman, 2000) (on September 14, 2023) for experimentally resolved structures of pathogen (viral or bacterial)-mammalian protein pairs. Only entries with a “Hetero 2-mer” (heterodimer) global stoichiometry and a resolution of 3.0 Å or lower were kept. The search results were also filtered to remove entries with chimeric proteins or experimental tags, entries with proteins without an assigned source organism, and entries where the source organisms in RCSB PDB differ from the source organisms of the associated UniProt entries. PDB sequence clusters at 70% sequence identity (from the September 13, 2023 weekly release of PDB) were then used to reduce redundancy in the dataset. The cluster ID for each protein pair was generated by combining the PDB sequence cluster IDs of the individual protein chains. For each unique pair cluster ID, only the entry with the largest number of modeled residues was selected. If multiple entries had the same number of modeled residues, only the one with the best resolution was kept. After filtering, the benchmark dataset had a total of 506 protein pairs, 250 viral-mammalian protein pairs and 256 bacterial-mammalian dimers. Structures for the first biological assembly were downloaded for all of the pairs, in PDB format. When the structures were only available in mmCIF format, we used the MAXIT suite from RCSB PDB to convert the files. The pdb_delinsertion tool from pdb-tools (Rodrigues et al., 2018) was used to remove insertion codes, renumbering the residues. Full sequences for each of the interacting proteins were obtained from UniProt (release 2023-05, downloaded on November 15, 2023) (The UniProt Consortium et al., 2023). When the provided Uniprot identifiers mapped to polyproteins, the name of the protein was used instead to find the sequence for the specific viral protein. For some proteins, such as antibodies and some viral proteins cleaved from polyproteins, the sequence provided by RCSB PDB was considered the “full” sequence. Since the experimental structures in PDB might not cover the full length of proteins and since partial sequences belonging to the same protein might have been assigned to different PDB sequence identity clusters, we clustered all of the full sequences using MMSeqs2 with a 70% sequence identity threshold and reassigned pair cluster IDs. The final benchmark dataset had a total of 456 protein pairs, 230 viral-mammalian protein pairs and 226 bacterial-mammalian dimers. The full protein sequences were used as input to AF-multimer.

### Host-pathogen protein pairs from published AP-MS experiments

High-confidence, experimentally identified host-pathogen interaction pairs were gathered from published AP-MS studies (*Chlamydia trachomatis* (Mirrashidi et al., 2015), *Mycobacterium tuberculosis* (Penn et al., 2018), Dengue and Zika (Shah et al., 2018), Ebola (Batra et al., 2018), HIV-1 (Jäger et al., 2012), HPV-31 (Eckhardt et al., 2018), KSHV (Davis et al., 2015), West Nile virus (WNV) (M. Li et al., 2019), SARS-CoV-1, MERS and SARS-CoV-2 (Gordon, Hiatt, et al., 2020; Gordon, Jang, et al., 2020), IAV (Haas et al., 2023), Hepatitis C virus (HCV) (Ramage et al., 2015), Hepatitis B virus (HBV) (Turnham et al., 2025) and CVB3 (Diep et al., 2019). The full sequences for the proteins involved in these pairs were obtained from the studies themselves or from UniProt (release 2023_05).

### HPIDB host-pathogen protein pairs

An additional set of host-pathogen PPIs was downloaded from HPIDB (R. Kumar & Nanduri, 2010), version 3.0. PPIs from X-ray crystallography experiments were removed, as well as any interactions already present in the AP-MS datasets described above. The dataset was also filtered by interaction type, and only physical associations, direct or not, were kept. A MIscore cutoff of 0.45 (Villaveces et al., 2015) was used to select high confidence interactions. We also removed any duplicate interactions, only keeping the row with the highest MIScore. We only kept human pathogens or pathogens that are commonly used as models for human pathogens in our final dataset. Sequences for the full proteins were downloaded from UniProt (2024-05 release), or from IntAct (Release 248) or NCBI RefSeq (Release 226), depending on the identifiers provided by HPIDB.

### Reducing protein pair redundancy

We clustered all sequences with a length of 10 amino acids or greater from the benchmark, AP-MS and HPIDB datasets using the easy-linclust workflow from MMSeqs2 to obtain cluster IDs for all of the protein pairs. The sequences were first clustered using the connected component algorithm (--cluster-mode 1), 80% coverage and a 70% sequence identity threshold. Pair cluster IDs were created by combining the sequence cluster IDs of the individual protein chains. The 70% sequence identity clusters were then used to remove redundant pairs from the benchmark dataset and to remove pairs in the AP-MS and HPIDB datasets that were highly similar to pairs in the benchmark dataset. The sequences were then clustered again using the connected component algorithm, a sequence coverage of 90% and a 95% sequence identity threshold. The resulting 95% sequence identity pair cluster IDs were used to reduce redundancy within the AP-MS and HPIDB datasets while allowing to maintain divergent protein sequences from different strains or highly related viruses.

### Human protein-protein interaction pairs

Host-pathogen PPI structures were compared with a previously published dataset of AF2 structures of human-human PPIs (Jänes et al., 2024). Only models with a pDockQ score of 0.49 or higher (“Medium” quality or better) were kept, for a total of 37,344 human PPI models.

### Structure prediction

Structures for the protein pairs from the benchmark and the AP-MS datasets were predicted using AF-multimer (version 2.3.1) (Evans et al., 2021). The use of templates was turned off when predicting structures for the benchmark datasets. ColabFold (version 1.5.5) (Mirdita et al., 2022) was used to predict structures for the interactions selected from HPIDB. AF-multimer was run with only one seed per model, producing a total of 5 predicted structures for each protein pair. All protein pairs were also modeled using AF3 (Abramson et al., 2024).

### Evaluating the quality of the predicted structures

Structures predicted for the benchmark datasets were evaluated by comparing them against the corresponding experimental structures and scoring them using two structural similarity metrics, the TM-score (Zhang & Skolnick, 2004), a measure of the global similarity between two structures, and the DockQ score (Basu & Wallner, 2016), which measures the quality of the interaction interface. TM-scores were calculated using MM-align (Mukherjee & Zhang, 2009) (2021/08/16 version). DockQ scores were calculated using DockQ version 1.0. A TM-score of 0.5 or greater indicates that two structures adopt similar folds (Xu & Zhang, 2010), while a DockQ score of 0.23 indicates that model is of at least “Acceptable” quality or better.

AP-MS and HPIDB structures were evaluated based on the confidence scores output by AlphaFold, mainly pTM, an estimate of the true TM-score, and ipTM, an estimate of the TM-score of the interface region only. The predicted local distance difference test (plDDT) was also taken into consideration when examining interfaces.

### Interface comparison

Interaction interfaces were defined by considering that any atom from one protein chain within an 8Å distance from any atom from the other protein chain is part of the interface. These interfaces are represented as lists of residue numbers.

The Jaccard index was used to quantify the similarity between different lists of interface residues belonging to the same protein extracted from different PPIs.

For each human protein (referred to as “bait”) in the AP-MS and HPIDB datasets, the interface residues targeted by the pathogen proteins that interact with it were extracted and clustered along with the interfaces extracted from human PPIs involving the same protein. The Jaccard distance was used to quantify the dissimilarity between interfaces. The interfaces were clustered using average linkage hierarchical clustering using scipy (version 1.10.1). The clustered interfaces were then represented as a heatmap of the “bait” interface residues targeted by the pathogen and human interactor proteins using seaborn (version 0.12.2).

Structural alignment of interfaces from human-pathogen and human-human protein pairs was performed using iAlign (Gao & Skolnick, 2010) (version 1.1), using the non-sequential alignment mode, a distance cutoff of 8Å, a minimum peptide length of 5 residues and a minimum interface size of 3 residues. The interfacial TM-score (iTM-score) was used as the similarity measure, and scores were normalized by the average length of both proteins.

The Dictionary of Secondary Structure in Proteins (DSSP) algorithm was used to assign secondary structure to the interface residues of pathogen and human protein interactors. The most frequently assigned DSSP code was determined for each interface and this was then used as additional information to confirm whether interface regions of pathogen and human interactors were structurally similar.

The pathogen interfaces were classified by similarity type based on manual inspection of the superposed structures of the pathogen-human and human-human protein pairs being compared.

### Gene set enrichment analysis

GO term enrichment analysis for human proteins involved in host-pathogen interactions with high confidence predicted structures was performed using clusterProfiler (version 4.14.6).

### Recombinant protein expression and purification

BL21(DE3) gold *E.coli* cells were transformed with constructs encoding the protein domains of interest, cloned into the pETM33 expression vector. Transformed cells were cultured in 2YT growth media (6 mg/mL peptone, 10 mg/mL yeast extract, 5 mg/mL NaCl) supplemented with 10% glycerol and 50 μg/mL kanamycin. Cultures were grown at 37 ℃ with shaking until reaching and optical density at 600 nm (OD_600_) of ∼0.6, at which point protein expression was induced with 0.1 mM isopropyl β-D-1-thiogalactopyranoside (IPTG) overnight at 18 °C.

Cells were harvested by centrifugation at 4 °C (4000 × g, 15 min) and resuspended in lysis buffer (PBS supplemented with 10% glycerol, 1% Triton X-100, 10 μg/mL DNase I, 5 mM MgCl_2_, 10 μg/mL of lysozyme, and cOmpleteTM EDTA-free Protease Inhibitor Cocktail (Hoffman-La Roche)). Cell lysis was performed by sonication (20s on, 15s off, 6 min), and the lysate was clarified by centrifugation (17000 x g, 45 min, at 4 °C). The supernatant was filtered through a 0.2 μm sterile PES filter and filtered lysates were incubated 1h with Cytiva Glutathione Sepharose™ 4 Fast Flow Media (Cytiva) and subsequently purified according to the manufacturer’s instructions. Briefly, unbound proteins were removed by washing three times with PBS, and bound GST-tagged proteins were eluted with 10 mM reduced glutathione (GTH) in PBS.

### SPOT arrays

Peptides covalently bound to a cellulose-ßalanine-membrane (PepSpots, JPT Peptide) were used to assess protein-peptide interactions. Membranes were first activated by incubation with 10 ml methanol for 5 min at room temperature (RT), followed by three 5-minute washes with 10 ml TBST (50 mM Tris, 150 mM NaCl, 0.05% Tween-20, pH 7.5). Membrane blocking was performed using 10 ml of 5% (w/v) skim milk powder (Merck Millipore, 115363) in TBST for at least 1h at RT. Blocked membranes were incubated overnight at 4 °C with the proteins of interest diluted in blocking buffer supplemented with 1mM dithiothreitol (DTT) under gentle rotation. The following day, membranes were briefly washed three times with TBST and subsequently incubated for 1h at 4 °C, with HRP-conjugated anti-GST antibody (Cytiva, RPN1236; 1:3000 dilution in blocking buffer) while rotating. After antibody incubation, membranes were washed three times for 2 min each in TBST, developed using Clarity Max Western ECL substrate (Biorad, 1705062) and imaged using the ChemiDoc Imaging system (Bio-Rad). Raw TIFF images were analyzed using ImageJ (Schneider et al., 2012) with signal intensities normalized to either the strongest signal or wild-type control (for alanine scanning arrays). Results are depicted as mean ± standard deviation (SD) from two replicates.

## Statistical analyses

The correlation (Spearman’s rank correlation coefficient) between the predicted model confidence scores (pTM, ipTM and pDockQ) and the TM-score or DockQ scores was calculated using scipy (version 1.10.1). Scores from different subsets of the benchmark dataset were compared using the Mann-Whitney U test, and scores for the same pairs modeled by AF-multimer and AF3 were compared using the Wilcoxon signed-rank test, both as implemented in the scipy package (version 1.10.1).

## Data and Code Availability

FASTA files and AF-multimer models for the benchmark, AP-MS and HPIDB host-pathogen pairs and PyMol session files for the models with the highest Jaccard index and iAlign scores are available from https://doi.org/10.5281/zenodo.15588018. The code used for this project can be found at https://github.com/dlrsb/host_pathogen_ppi_struct_pred.

## Supporting information

SuppTables

## Acknowledgments

This research was funded by a grant from the NIH (PTE Federal Award No: 1U19AI171110-01 Subaward No:13668sc). MJA has received funding from “la Caixa” Foundation and FCT, I.P. under the project code [LCF/PR/HR22/00722]. Structure prediction and interface comparison analyses were carried out on the ETH Zürich Euler cluster. PB was supported by the Helmut Horten Stiftung and the ETH Zurich Foundation.

## Supplementary Materials Supplementary Tables

**Table S1 -** Benchmark dataset of viral-mammalian and bacterial-mammalian protein pairs selected from PDB used to evaluate the performance of AF-multimer and AF3.

**Table S2 -** Model quality scores for the AF-multimer structures predicted for the benchmark pairs.

**Table S3 -** Model quality scores for the AF3 structures predicted for the benchmark pairs.

**Table S4 -** Dataset of human-pathogen protein pairs selected from AP-MS studies and HPIDB.

**Table S5 -** Predicted model confidence scores for the AF-multimer structures predicted for the AP-MS and HPIDB pairs.

**Table S6 -** Predicted model confidence scores for the AF3 structures predicted for the AP-MS and HPIDB pairs.

**Table S7 -** Human-human protein-protein interaction pairs from (Jänes et al., 2024) used in this study (structures with a pDockQ score of 0.49 or higher).

**Table S8 -** Pathogen-human and human-human protein-protein interaction pairs considered for interface comparison.

**Table S9 -** Number of pathogen proteins per human (“bait”) protein.

**Table S10 -** Interface similarity scores calculated between human-pathogen interfaces sharing the same human (“bait”) protein.

**Table S11 -** Interface similarity scores calculated between human-pathogen and human-human interfaces sharing the same human (“bait”) protein.

**Table S12 -** Pathogen interfaces classified by the type of similarity they share with human interfaces. Interfaces were classified considering the human-human pairs that scored the highest in terms of Jaccard index and iAlign iTM-score.

**Table S13 -** Mimicry of short linear motifs detected among pathogen proteins with linear/less structured interface regions.

**Table S14 -** Pathogen proteins targeting interfaces not targeted by human proteins.

---

**Figure S1.**
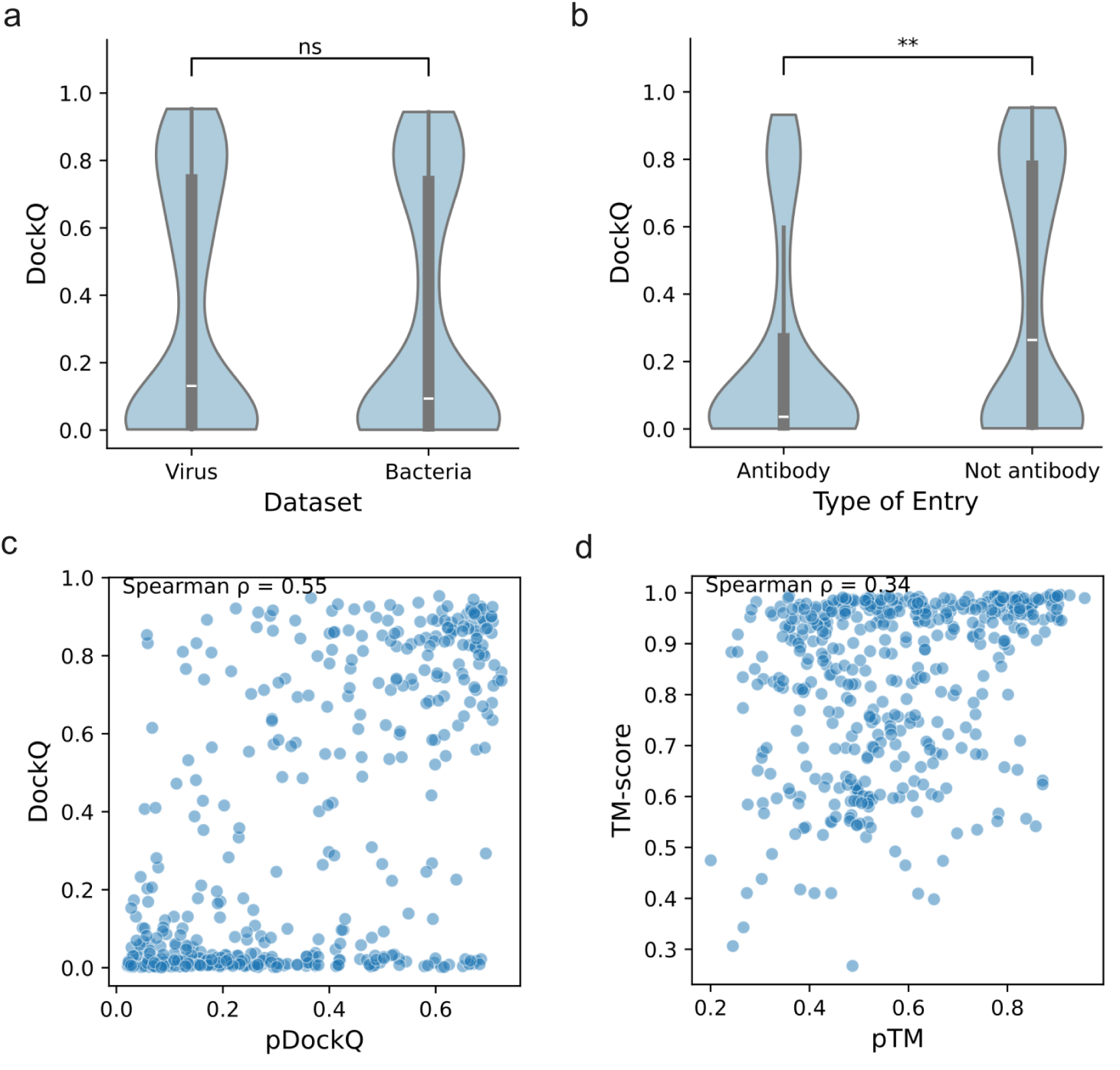
Results for the benchmark host-pathogen protein pair structures predicted using AF-multimer. a) Comparison between the viral-mammalian and bacteria-mammalian protein pairs in the benchmark dataset, in terms of DockQ scores. DockQ scores are not significantly different between the viral and bacterial protein pairs (Mann-Whitney U test p-value = 0.57). b) Comparison between protein pairs in which one of the interactors is an antibody (N = 119) and pairs without antibodies (N=333). DockQ scores are significantly lower for protein pairs involving antibodies (Mann-Whitney U test p-value = 0.001). c) Correlation between pDockQ (as defined in (Burke et al., 2023)) and DockQ scores for AF-multimer structures (Spearman’s ρ = 0.55). d) Correlation between pTM and TM-score values for AF-multimer structures (Spearman’s ρ = 0.34).

**Figure S2.**
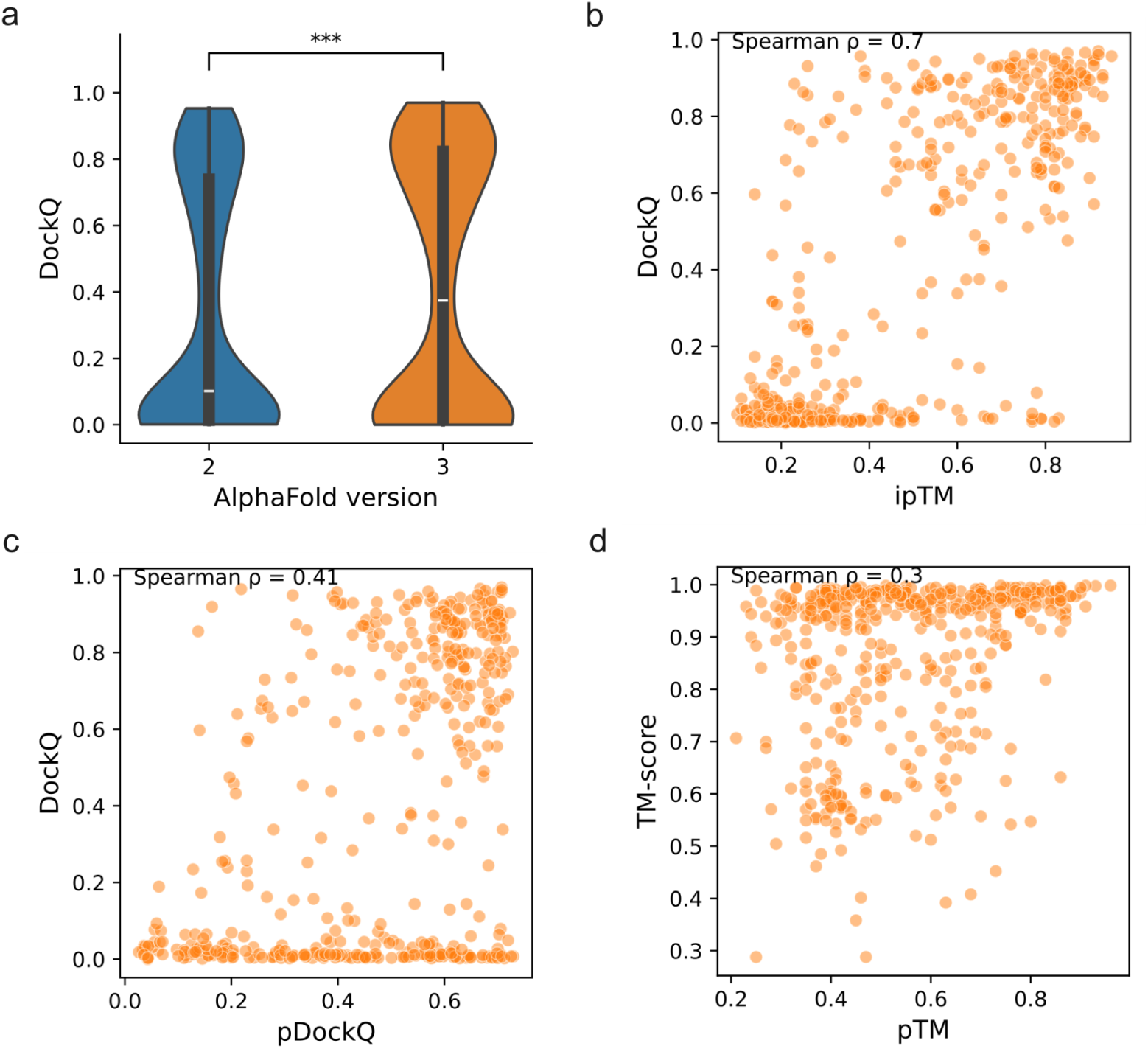
Results for the benchmark host-pathogen protein pair structures predicted with AF3. a) Comparison of DockQ scores for structures predicted with AF-multimer and AF3. AF3 structures for the benchmark protein pairs have significantly higher DockQ scores than the AF-multimer structures (Wilcoxon signed-rank test p-value = 1.36 x 10^-5^). b) Correlation between ipTM scores and DockQ scores for AF3 structures (Spearman’s ρ = 0.70). c) Correlation between pDockQ (as defined in (Burke et al., 2023)) scores and DockQ scores for AF3 structures (Spearman’s ρ = 0.41). d) Correlation between pTM and TM-score values for AF3 structures (Spearman’s ρ = 0.30).

**Figure S3.**
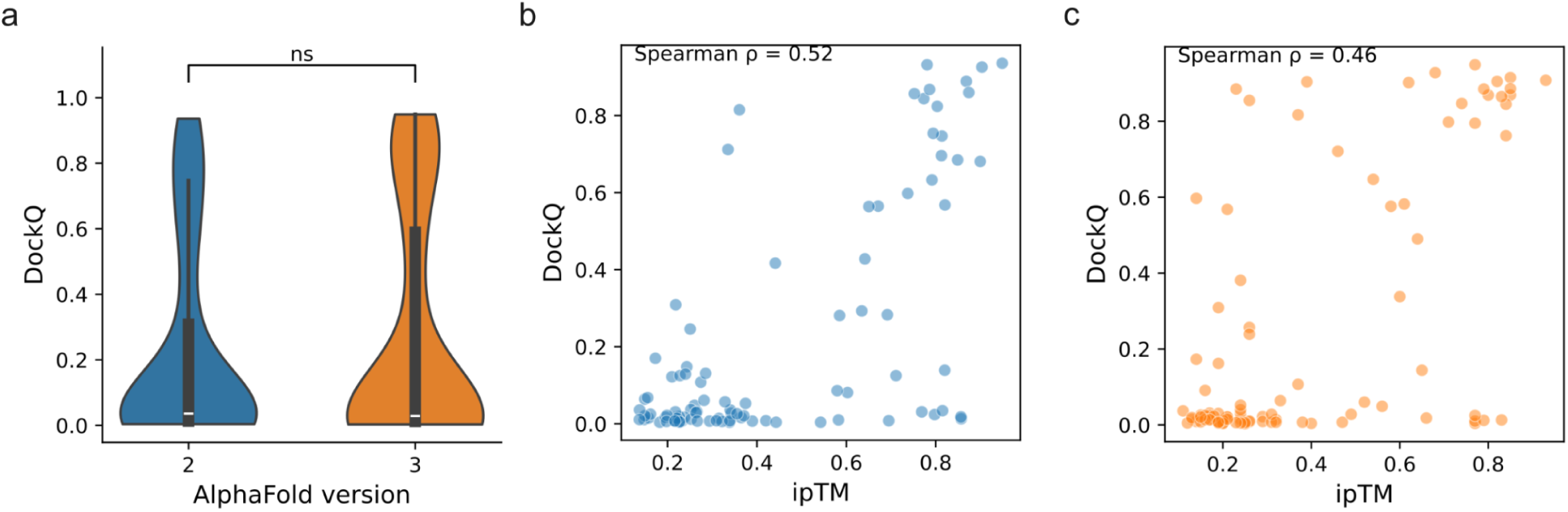
Results for the benchmark host-pathogen protein pairs that were released after the AF-multimer v. 2.3.0 and AF3 training cutoff date (30 September 2021). a) Comparison of DockQ scores for structures predicted with AF-multimer and AF3. DockQ scores for the protein pairs released after the training cutoff are not significantly different between AF-multimer and AF3 (Wilcoxon signed-rank test p-value = 0.55). b) Correlation between ipTM scores and DockQ scores for structures predicted with AF-multimer, when only considering the benchmark pairs released after the training cutoff date (Spearman’s ρ = 0.52). c) Correlation between ipTM scores and DockQ scores for structures predicted with AF3, when only considering the benchmark pairs released after the training cutoff date (Spearman’s ρ = 0.46).

**Figure S4.**
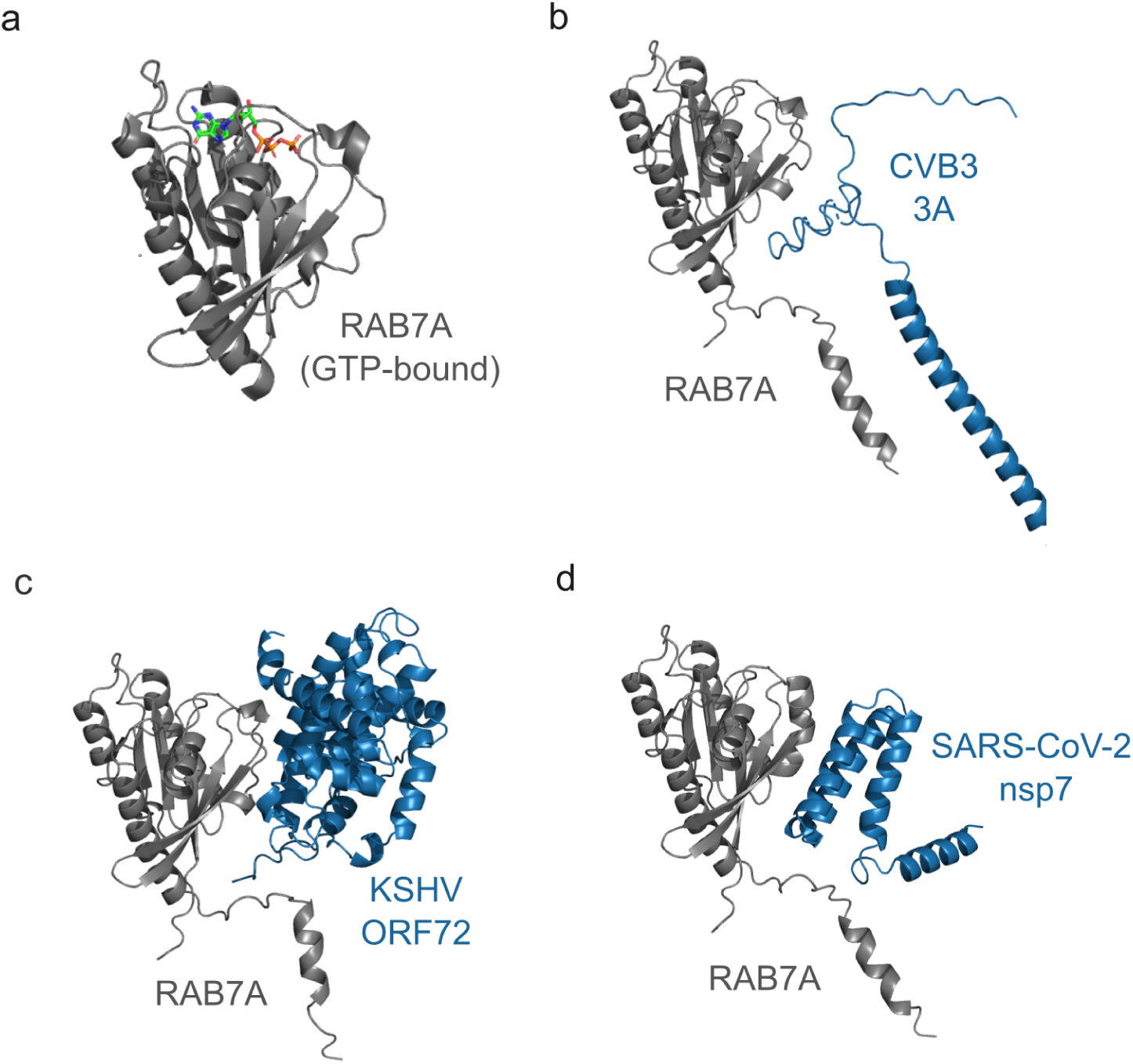
Comparison between an experimental structure of GTP-bound RAB7A (in gray) and AF-multimer predicted structures for RAB7A interacting with 3 different viral proteins (in blue). a) Crystal structure of RAB7A binding GTP (PDB 1T91 (M. Wu et al., 2005)). b) Predicted structure for the interaction between RAB7A and CVB3 3A. c) Predicted structure for the interaction between RAB7A and KSHV ORF72. d) Predicted structure for the interaction between RAB7A and SARS-CoV-2 nsp7.

**Figure S5.**
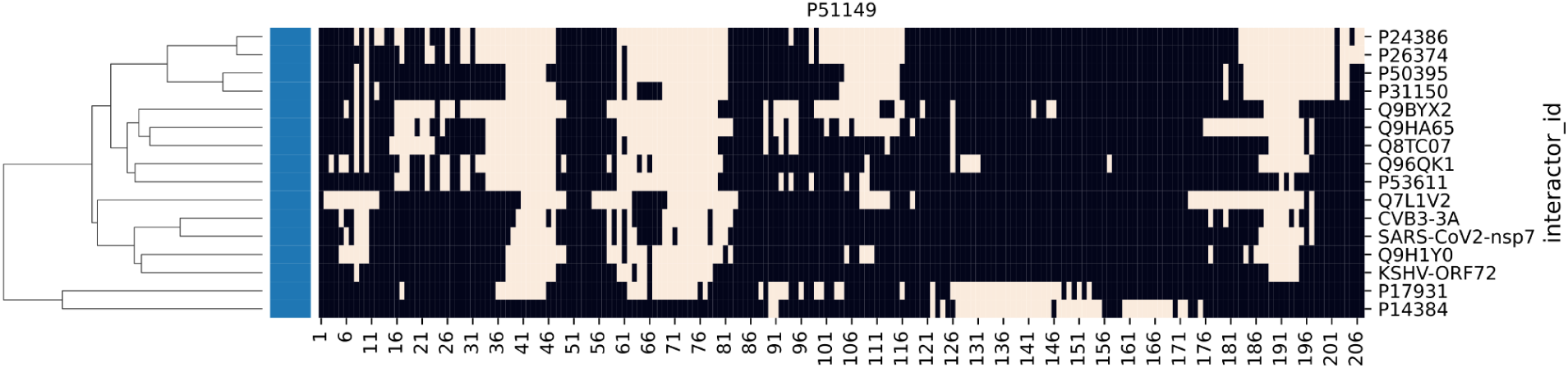
Residues from Ras-related protein Rab-7a (RAB7A, P51149) targeted by various human and pathogen proteins (interactor_id). Targeted residues are shown in beige, and the remaining residues are shown in black. The targeted interface residues were clustered based on the Jaccard distance.

**Figure S6.**
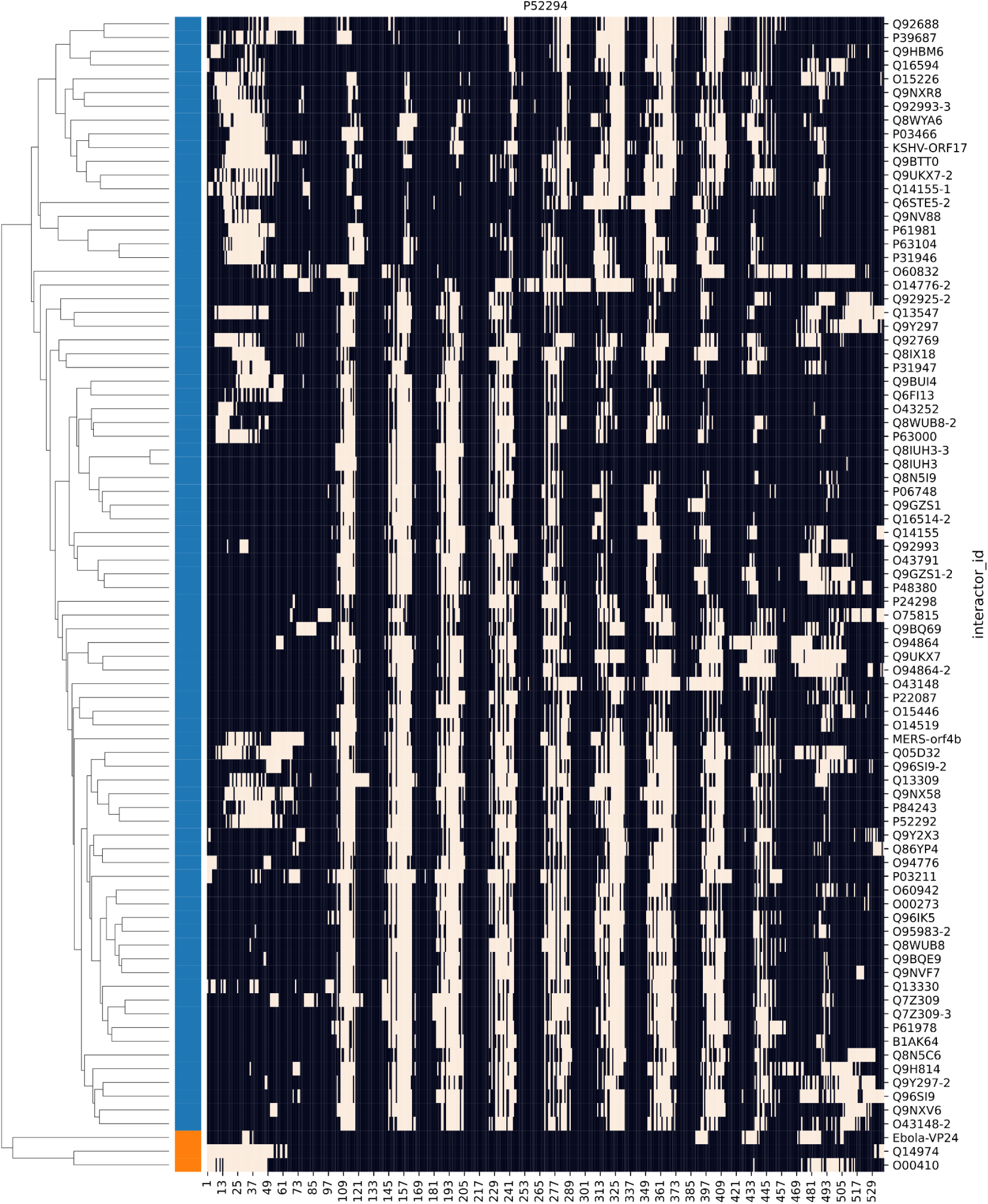
Residues from Importin subunit alpha-5 (KPNA1, Uniprot ID P52294) targeted by various human and pathogen proteins (interactor_id). Targeted residues are shown in beige, and the remaining residues are shown in black. The targeted interface residues were clustered based on the Jaccard distance.

**Figure S7.**
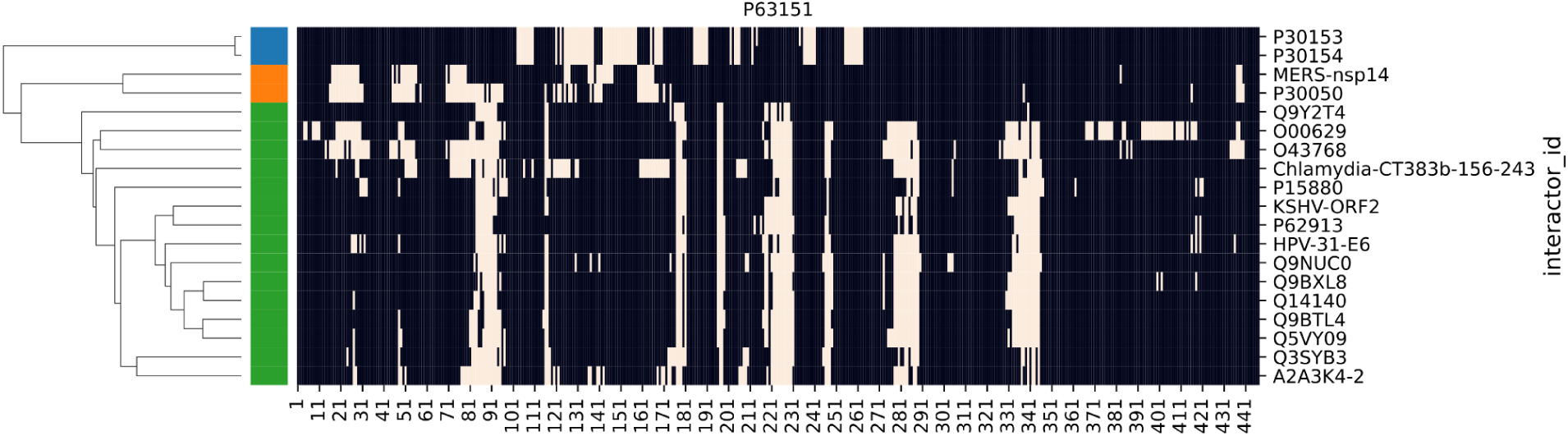
Residues from Serine/threonine-protein phosphatase 2A 55 kDa regulatory subunit B alpha isoform (PPP2R2A, Uniprot ID P63151) targeted by various human and pathogen proteins (interactor_id). Targeted residues are shown in beige, and the remaining residues are shown in black. The targeted interface residues were clustered based on the Jaccard distance.

**Figure S8.**
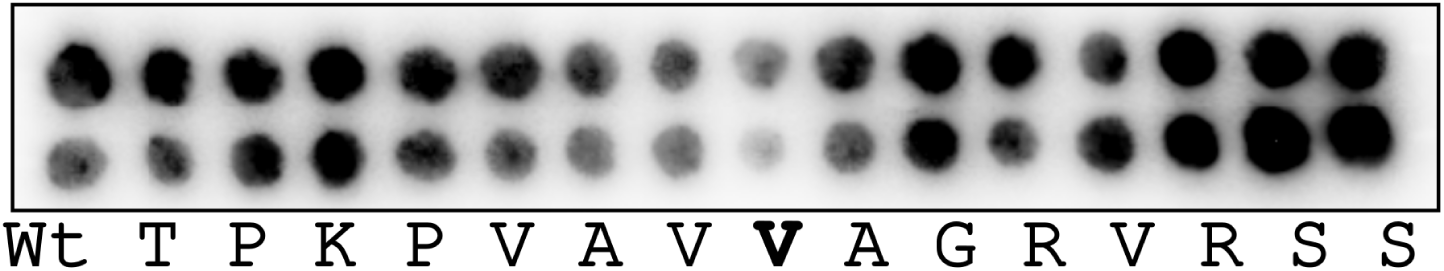
Alanine scanning experiment for the interaction between SIAH1 and a peptide from the predicted interface residues in KSHV ORF45 (TPKPVAVVAGRVRSS).

